## Supplemental Figures 1-3 for "Antibody Profiles Elicited by Potent and Subpotent Whole Cell Pertussis Vaccines in Mice"

**Tables S1-S9. Provided as Excel File**

**Reactive *B. pertussis* antigens recognized by wP immune sera at all stress temperatures relative to sham vaccinated animals**

Figure S1

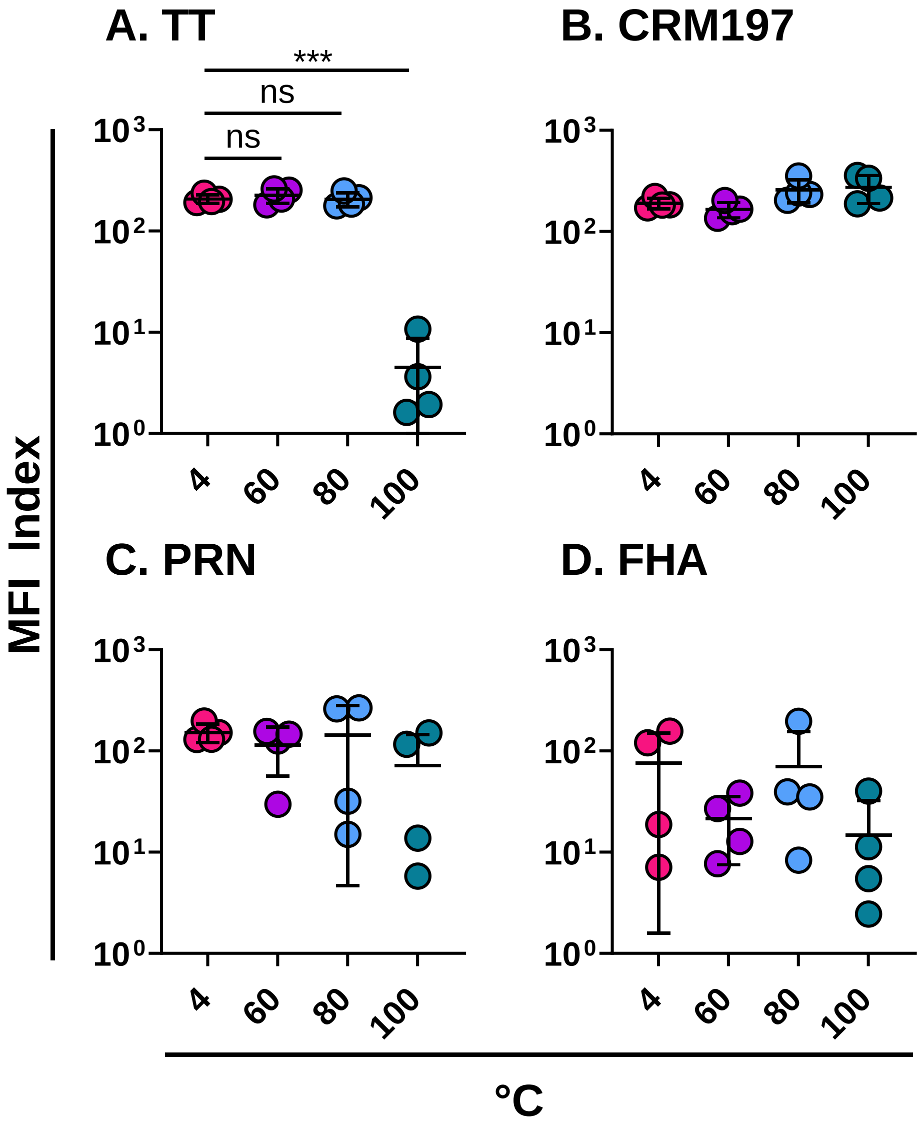


**Figure S1. Antigen-specific responses to DTwP$_{4, 60, 80 ,100.}$** Mice (n=4 per group) were intraperitoneally vaccinated with DTwP stressed at different temperatures on days 0 and 21 with 1.2 OU/mouse. Sera was collected on day 30 pre-challenge and was analyzed by MIA to determine antigen-specific titers.

### Figure S2

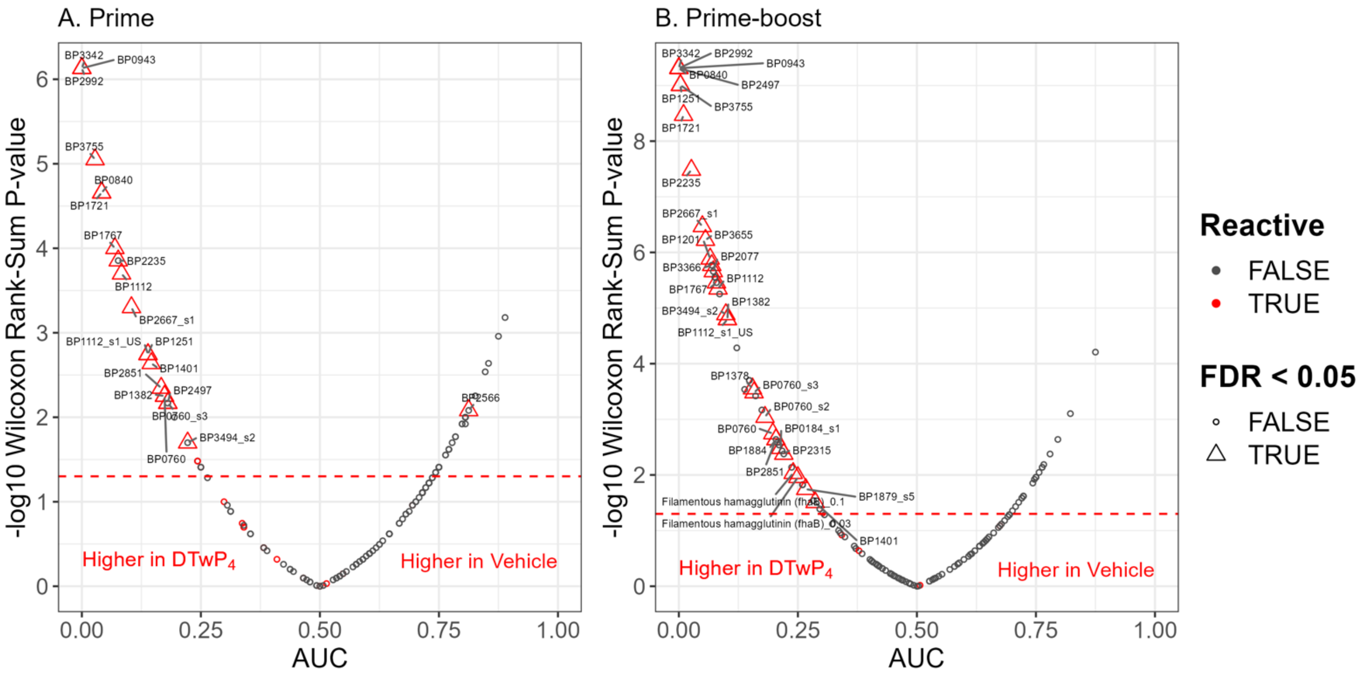


**Figure S2. Antigens specifically associated with $DTwP_4$ vaccination.** Serum samples from BALB/c mice (n=51) vaccinated with $DTwP_4$ on day 0 (prime) or day 0 and 21 (boost); alongside sham vaccinated mice (n=44) were subjected to the *B. pertussis* proteome array. The identified immunoreactive IVTT *B. pertussis* antigens were rank ordered based on AUC and statistical significance.

#### Figure S3

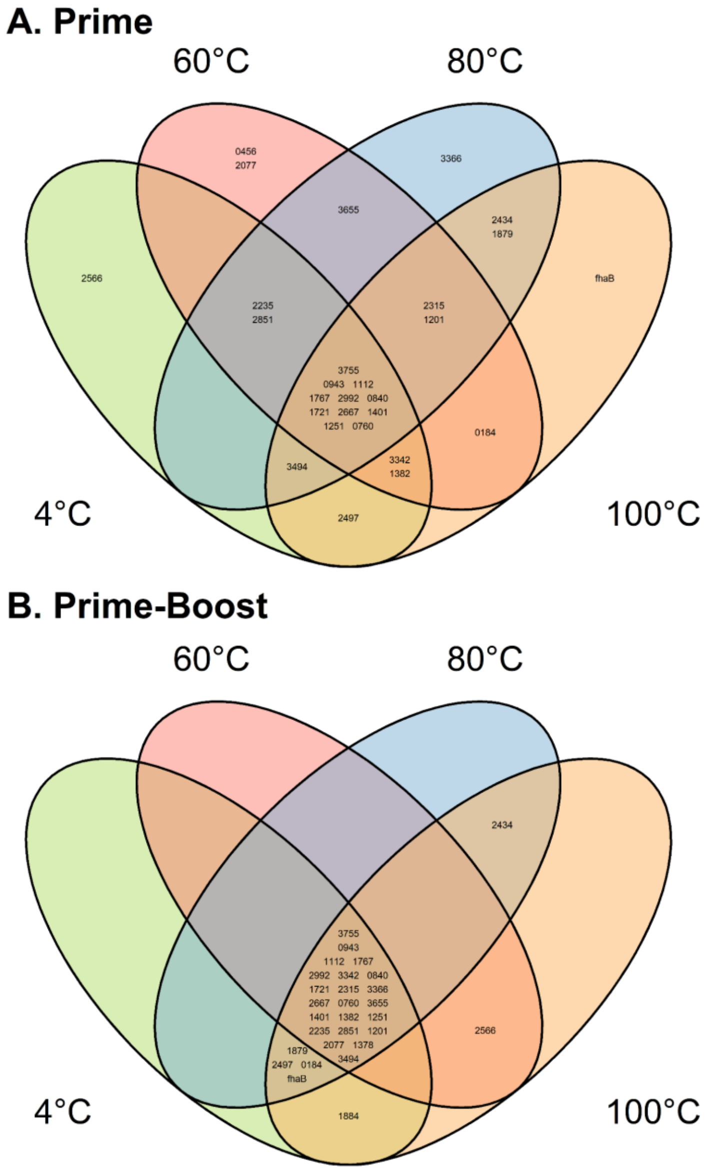


**Figure S3. Venn diagram showing overlaps between differential reactive antigens across all stress temperatures.** To identify unique serological signatures, differential reactive antigens in the serum samples from prime (d0) and boost (d0 and d30) $DTwP_{4,\ 60,\ 80\ ,100}$ vaccinated animals were compared for overlaps at all stress temperatures.
